# Reassortment, positive selection, and the inter-segmental patterns of divergence and polymorphism in influenza virus H3N2

**DOI:** 10.1101/360941

**Authors:** Kangchon Kim, Yeongseon Park, Yuseob Kim

**Author notes:** Corresponding author: Yuseob Kim, Department of Life Science, Ewha Womans University, Ewhayeodae-gil 52, Seodaemun-gu, Seoul, Korea 03760.

## Abstract

Reassortment in viruses with segmented genome is a major evolutionary process for their genetic diversity and adaptation. It is also crucial in generating different levels of sequence polymorphism among segments when positive selection occurs at different rates on them. Previous studies have detected intra-subtype reassortment events in human influenza H3N2 by between-segment incongruity in phylogenetic tree topology. Here, we quantitatively estimate the reassortment rate, probability that a pair of segments in a viral lineage become separated in a unit time, between hemmaglutinin (HA) and four non-antigenic segments (PB2, PB1, PA and NP) in human influenza virus H3N2. Using statistics that measure incongruity in tree topology or linkage disequilibrium between segments and performing simulations that are constrained to reproduce the various patterns of H3N2 molecular evolution, we infer that reassortment rate ranges between 0.001 and 0.01 assuming one generation to be 1/80 year. However, we find that a higher rate of reassortment is required to generate the observed pattern of ~40% less synonymous sequence polymorphism on HA relative to other non-HA segments, which results from recurrent selective sweeps by antigenic variants on the HA segment. Here, synonymous diversity was compared after correcting for difference in inferred mutation rates among segments, which we found significant. We also explored analytic approximations for inter-segmental difference in sequence diversity for a given reassortment rate to understand the underlying dynamics of recurrent positive selection. It is suggested that the effects of clonal interference and potentially demography-dependent rate of reassortment in the process of recurrent selective sweeps must be considered to fully explain the genomic pattern of diversity in H3N2 viruses.

---

The evolution of influenza virus has been one of major long-standing subjects of modern biological researches, owing to its significant impact not only on human public health but also on the study of adaptive molecular evolution (Fitch *et al*. 1991; Yang 2000; Nelson and Holmes 2007). Studies focused on the sequence evolution of hemagglutinin (HA) and neuraminidase (NA) gene segments because HA and NA proteins are recognized as antigens by host adaptive immune system. Rapid amino acid substitutions at their epitope sites cause “antigenic drift” that forces updates in flu vaccines. The occurrence of positive selection on these sites was confirmed by various evidences and is now generally believed to drive the evolutionary dynamics of influenza viruses. However, given that the complex seasonal dynamics of viral populations has its own effect on the temporal patterns of sequence diversity or polymophism and that multiple antigenic sites, together with other functional sites on a non-recombining segment (“complete linkage” within a segment), undergo correlated evolutionary changes, it is not easy to identify and analyze selection from the observation of viral sequence evolution (Illingworth and Mustonen 2012; Strelkowa and Lassig 2012; Kim and Kim 2015).

The correct evolutionary model of influenza virus must predict the observed patterns of sequence diversity as accurately as possible. The observed patterns include the characteristic “cactus-like” genealogical trees for individual viral segments (Buonagurio *et al*. 1986; Ferguson *et al*. 2003; Bedford *et al*. 2011), the ratio of nonsynonymous versus synonymous sequence changes at epitope and non-epitope sites (Ina and Gojobori 1994), the relative distribution of amino acid substitutions in external versus internal branches of trees (Fitch *et al*. 1997; Pybus *et al*. 2007), the absolute level and geographic differentiation of sequence diversity (Bedford *et al*. 2010), and the temporal correlation of variants’ fixation events (Strelkowa and Lassig 2012; Kim and Kim 2015). These patterns were mostly observed in the HA segment and therefore models were developed mainly to explain the evolution of HA gene. However, the evolutionary change of HA is not independent of other segments. Unless viruses co-infect hosts very frequently and exchange segments with each other, thus resulting in very frequent reassortment, different segments in a viral lineage are not separated during most of the infectious cycles. Such correlated inheritance, or genetic linkage across segments, means that evolutionary events on one segment can affect the dynamics of others. The pattern of genetic variation observed in HA is therefore expected to be shaped by the fitness effects of variants not only on HA but also on other segments.

The HA segment of H3N2 viruses exhibit lower level of genetic diversity, measured in mean time to coalescence, than other segments (Rambaut *et al*. 2008). This is explained by recurrent positive selection occurring at far higher rate on the HA segment than other segments. Selective sweeps driven by antigenic variants on HA therefore cause the greatest reduction in polymorphism at linked sites on the same segment but less severe reduction at other segments due to occasional events of reassortment that break down the hitchhiking effect (Maynard Smith and Haigh 1974). Relative diversity between segments is therefore informative for adaptive evolution in the HA gene. In addition, negative (or purifying) selection against deleterious mutations cause reduction in polymorphism, an effect termed background selection (Charlesworth *et al*. 1993). This variation-reducing effect is also greatest on completely linked sites and diminishes as linkage becomes weaker. Since negative selection must be operating in all genes of influenza virus to maintain their functions, genetic diversity at HA must be affected not only by negative selection on the same segment but also that on all the other segments, unless reassortment is very frequent relative to the strength of negative selection.

Therefore, the evolutionary model of positive and negative selection should be tested against the inter-segmental levels and patterns of sequence polymorphism. However, a crucial parameter in such a model with multiple viral segment, the rate of reassortment between segments, is not well known. Reassortment in segmented RNA virus, effectively equivalent to meiotic recombination in most eukaryotes, plays a critical role in their evolution. To date, eleven families of RNA virus are known to have segmented genome (McDonald *et al*. 2016). Among these, reassortment in influenza virus has been most intensively studied. Through this process, influenza viruses can acquire novel variation that confers resistance to antivirals (Simonsen *et al*. 2007). Intrasubtype reassortments also drive adaptive amino acid replacements. Past pandemics have been attributed to the result of reassortment between different influenza subtypes (Nelson and Holmes 2007). Therefore, detecting and understanding reassortment has been of great public health interest.

Numerous studies have detected reassortment from serially sampled influenza virus sequences. Reassortments were observed within and between subtypes of human influenza A (Holmes *et al*. 2005; Schweiger *et al*. 2006; Lycett *et al*. 2012; Lu *et al*. 2014; Westgeest *et al*. 2014; Pinsent *et al*. 2015; Berry *et al*. 2016; Villa and LÄssig 2017), and between lineages of influenza B virus (Dudas *et al*. 2014). While it can be identified manually by comparing phylogeny between segments, for comprehensive analysis and identification computational detection algorithms were suggested. Most widely used method detects a clade that occupies a position in a phylogenetic tree constructed for one segment is located on a different position in the corresponding tree for a different segment (Nagarajan and Kingsford 2010). Such a clade thus represents a reassortant. Other methods are not dependent on phylogeny. Rabadan *et al*. (2008) identified the presence of reassortment when mean sequence difference between two taxa is highly variable for different segments. However, this approach overlooked the possibility that different segments may have different sequence diversity not due to reassortment but due to segment-dependent effective population sizes.

Despite these sophisticated methods for identifying reassortment, rare attempt has been made to estimate how frequently it occurs during viral reproduction, particularly in comparison to the rates of mutation and coalescence. The rate of reassortment per unit time (∆*t*) can be defined as a probability that a pair of segments in a given individual virus at time *t* come from different parental viruses that existed at time *t* - ∆*t*. If the reproduction of viruses can be approximated in a discrete-time process, a natural choice for the unit time above can be the average length of a single host infection cycle, which we arbitrarily define to be one “generation” (Kim and Kim 2016). In previous studies, reassortment rate was often estimated as the number of detected events divided by years or the number of synonymous changes on the tree (Villa and LÄssig 2017). This quantity may be a lower bound of the actual rate since only those events leaving sufficiently conspicuous inter-segmental incongruence in phylogenies are counted. In this study, we perform a quantitative analysis of reassortment rate in influenza H3N2, using summary statistics (“metrics”) that measure either incongruity in tree topology or linkage disequilibrium. We conduct simulations of viral sequence evolution under four different models, including recurrent positive selection with and without complex demography, that are however constrained to replicate the key patterns of H3N2 sequence variation. Then, the range of reassortment rate that reproduces the observed values of these metrics as well as the ratio of sequence diversity at HA versus non-HA segments will be identified. We also seek analytic approximations for the effect of recurrent positive selection with varying reassortment rate and other theoretical explanations to understand inter-segmental variation in the level of polymorphism observed in the actual and simulated sequences.

## MATERIALS and METHODS

### Sequence data

Genome sets of human influenza A/H3N2 sequences were downloaded from Influenza Virus Genome Set of National Center for Biotechnology Information (NCBI). A genome set is defined as the sequences of viral segments from a single virus isolate. In this study, we use sequences of HA, PB2, PB1, PA and NP segments. Outlier sequences (different from other sequences in the same year at more than 100 sites) and sequences containing symbols other than A, C, G and T were discarded.

### Statistics for inter-segmental genetic correlation

Robinson-Foulds distance (RFD; Robinson and Foulds 1981) was calculated between evolutionary trees from different viral segments to quantify incongruence between their topologies. Neighbor-joining trees constructed from individual segments were fed into TreeDist in PAUP 4 test version (Swofford 2003).

The standard measure of linkage disequilibrium (LD) for a pair of bi-allelic sites, ρ^2^, is calculated as

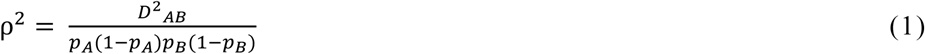

where *p_A_* is the frequency of allele *A* at locus 1 and *p_B_* is the frequency of allele *B* at locus 2 and *D_AB_* is *p_AB_* - *p_A_p_B_* (Hill and Robertson 1968). To quantify LD between segments, ρ^2^ is calculated for each pair of sites, one on segment 1 and another on segment 2. The average of all such pairs is given by 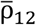. We define a metric that quantifies between-segment LD relative to within-segment LD as

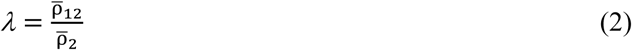

where 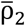 is the mean of ρ^2^ between all pairs of sites within segment 2, which is either a non-HA segment in actual data or a segment that evolves without positive selection in simulation (see below).

Topology based linkage disequilibrium (TBLD), proposed recently by Wirtz *et al*. (2018) as an improvement over conventional SNP-based LD, is obtained by grouping sequences of a segment into two alleles defined by tree topology, as illustrated in Figure 1. Then, ρ^2^ is calculated between segments using the frequencies of such topology-based alleles. For this analysis, neighbor-joining trees constructed above for Robinson-Foulds metric were used again.

**Figure 1.**
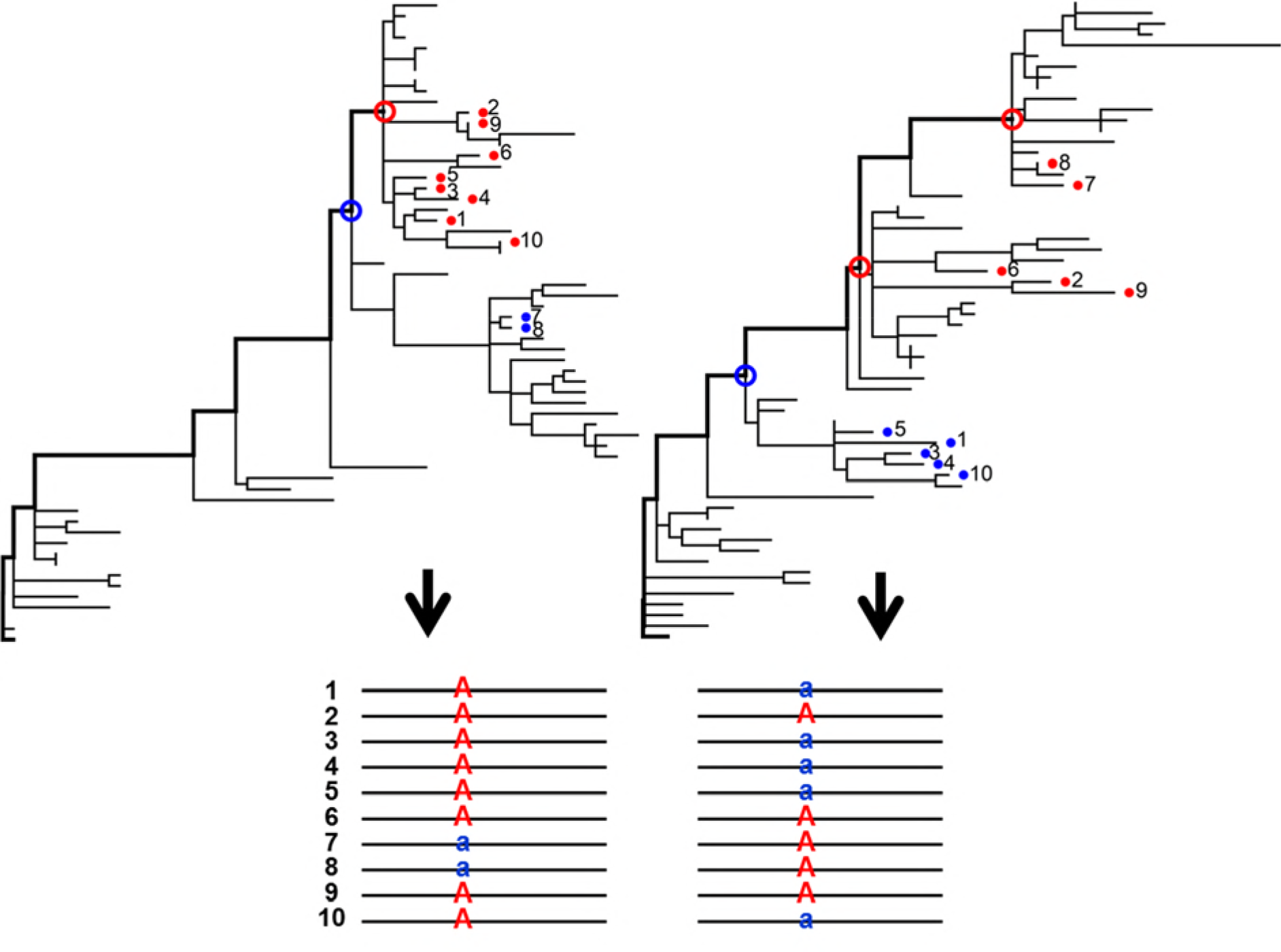
Figure legend: Topology-based linkage disequilibrium (TBLD) method. (A) Phylogenies are constructed from two segments. From a phylogeny of a segment, each taxon within a 6-month time window (numbered from 1 to 10) is traced back to the tree trunk (thicker line) and is mapped to the “first node” encountering the tree trunk on its way (empty circles). Then we grouped taxa into two according to their “first nodes”: the first group (blue filled circles) consists of taxa mapped to the most ancestral first node (blue empty circle) and the second group (red filled circles) consists of taxa mapped to the other first nodes (red empty circles). (B) Taxa are labeled according to their group so that *r*^2^ is calculated to quantify TBLD.

The above metrics were calculated for 30 genomic sets (from either actual H3N2 or simulated population) randomly sampled within each 6-month time window. For H3N2 data, sequences from different regions (Asia, Europe, North America, South America, Oceania and others) were sampled proportionally to the number of sequences in the database. For a given metric, the average value over time windows, from year 2007 to 2016 for H3N2 data or over 10 simulation years, was obtained.

### Simulation

We conducted the individual-based simulation of virus evolution in a procedure described in Kim and Kim (2016) with modification. In this study, a virus consists of two segments, each containing 1,000 bi-allelic sites. Segment 1 is modeled after the HA1 segment and have 770 “nonsynonymous” sites including *L*_b_ “epitope” sites where beneficial mutations occur to increase viral fitness by *s*. On the other hand, segment 2 does not have epitope sites. Other sites on segment 1 or 2 are either under negative selection (770 – *L*_b_ nonsynonymous sites that mutate to deleterious alleles with selection coefficient *s*_d_) or under neutral evolution (*L*_s_ = 230 “synonymous” sites). The population evolves in discrete generations, with one generation corresponding to 1/80 year. Mutation rate per site per year is given by *μ* = 8.0 ×10^−3^ (10^−4^ per generation) which is approximately the estimate of per-nucleotide mutation rate in H3N2 viruses.

As described in Kim and Kim (2016), after steps of migration and mutation, a Poisson number of progenies are produced per a parental copy of virus as a function of its absolute fitness, which is obtained by multiplying its relative fitness (after combining effects of all beneficial and deleterious mutations) by the ratio of population size to carrying capacity (*K*) at each generation. Let *N* be the number of viruses after these steps of reproduction. Then, we randomly select two viruses that exchange their segments with probability 0.5. This step is repeated *Nr* times. Therefore, the rate of reassortment per viral lineage per generation is *r*.

Four evolutionary models are considered. First, in model 1, both segments are subjected only to genetic drift in a near constant-sized population (thus *s* = *s*_d_ = 0). Here, carrying capacity (*K* = 140 ≈ *N*) was given to yield π_1_ = 0.027, the mean pairwise sequence difference per site in segment 1, which corresponds to the observed synonymous diversity in the HA segment of H3N2 population. We also consider models with recurrent positive selection without (model 2; *s*_d_ = 0) or with (model 3; *s*_d_ > 0) negative selection at other nonsynonymous sites on both segments. The strength of positive selection, *s*, is set either 0.05 or 0.1, as we previously estimated *s* to range between 0.05 and 0.11 by examining how rapidly the frequencies of known antigenic-cluster-changing variants (Koel *et al*. 2013) increase over time (Kim and Kim 2016). In all models with selection, *N* was adjusted to yield π_1_ very close to 0.027. Other evolutionary parameters relevant for segment 1 in model 2 and model 3 are identical to those of Model A and Model B1 (*L* = 1,000), respectively, in Kim and Kim (2016). Finally, we examine the model of positive selection only, but together with metapopulation dynamics (model 4; *s* = 0.1 and *s*_d_ = 0). This model uses the same parameter values as in the Model C3a (constant carrying capacity in tropical region) of Kim and Kim (2016). Briefly, the metapopulation consists of ten local demes, each of which sends migrants in proportion to its size to other demes. Five and two demes are colonized in “winter” and “summer”, respectively, and go extinct in the next season. A remaining deme, modeling a tropical population with continuous influenza epidemics, however is maintained without extinction.

### Synonymous diversity and divergence

To obtain synonymous diversity π for each segment, mean pairwise synonymous difference among sequences sampled within a 6-month window was calculated according to Nei-Gojobori method (Nei and Gojobori 1986). Then, the average over 20 years from 1997 to 2016 (40 time windows) was obtained. Synonymous diversity corrected for mutation rate, π*, is obtained by dividing π by the synonymous divergence of corresponding segment from 1997 to 2016 (see below). Tajima’s D (Tajima 1989) was also calculated for 30 sequences in each of the above 6-month windows and the average over windows was obtained.

To estimate synonymous divergence, which is the number of nucleotide substitutions per synonymous sites, we reconstructed phylogeny for each PB2, PB1, PA, HA and NP segment. We tracked ancestral sequences at all internal nodes of phylogeny on a path starting from tree root to sequences sampled at each year and counted the cumulative number of synonymous changes on the path. Measuring cumulative divergence along the phylogeny, rather than just calculating synonymous differences between two terminal years of sampling, prevents multiple nonsynonymous changes at one site being counted as a synonymous change, especially in HA1 domain, or multiple synonymous changes at one sites being counted as a smaller number of changes. Neighbor-joining trees were reconstructed using PAUP with 30 sub-sampled sequences per year from 1973 to 2016 from all available sequences from Genbank. The trees were rooted to the common ancestor of sequences collected in 1973 because, across all segments, these sequences exhibit very little diversity and therefore their common ancestor is confidently dated to the same year. Internal node states were inferred to track synonymous changes along the branch using ACCTRAN method in PAUP. For this analysis, we used either four-fold synonymous sites or all synonymous sites. To obtain synonymous divergence for the latter, we used the Nei-Gojobori method.

To test whether the rate of sequence divergence at one segment is significantly different from that of another segment, bootstrap test was performed according to (Hall and Wilson 1991). Let *d_X_* and *d_Y_* be the divergence of segment *X* and *Y*. Then we define 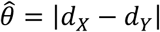 and test if it is significantly greater than zero. As the distribution of 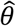 under the null hypothesis (*d_X_* = *d_Y_*) can be approximated by the distribution of 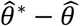 where 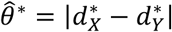 is a bootstrap value of 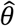, the P-value is approximately the proportion of bootstrap samples that satisfy 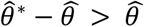. For each pair of segments, 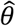 was obtained from divergences from 1973 to 2016 calculated by the above method. A pseudo data set is prepared by randomly sampling triplet-codon columns in the alignment of a given segment with replacement until it has the same number of codons as the original sequence. For bootstrap test, 1000 pseudo data sets for each segment pair were generated.

### Data availability statement

The authors affirm that all data necessary for confirming the conclusions of this article are represented fully within the article and its tables and figures

## RESULTS

### The estimation of inter-segmental reassortment rate in influenza virus H3N2

Population genetic processes at different segments become uncorrelated as reassortment occurs. Therefore, we attempted to infer reassortment rate in H3N2 viruses using multiple summary statistics that measure correlation in the patterns of sequence diversity across segments. One metric we use is Robinson-Foulds distance (RFD) between evolutionary trees, each of which is constructed from sequences of one particular segment (Robinson and Foulds 1981). As a measure of tree incongruity, RFD is expected to be positively correlated with reassortment rate. On the other hand, linkage disequilibrium (LD) between polymorphism on different segments is expected to decay with an increasing rate of reassortment. We consider two summary statistics (metrics) of inter-segmental LD, λ and TBLD (see Methods).

To investigate whether these metrics are sufficiently informative and robust for inferring reassortment rate, we performed simulation of virus population in which two segments (one modeling the HA segment and the other a non-HA segment) are undergoing varying rates of reassortment (*r* = 0 to 10^−2^). The relationship between a given metric and reassortment rate may depend on the pattern of sequence diversity, which is determined by how viruses evolve. We therefore simulated virus population under four distinct population genetic models: simple neutral evolution (model 1), recurrent positive selection (selective sweeps; model 2), recurrent positive and negative selection (model 3), and positive selection under complex demographic dynamics (model 4). Parameters of each evolutionary model were adjusted to yield a constant level of synonymous sequence diversity (or effective population size *N*_e_ ≈ 140) and constant rate of adaptive substitutions (*k* ≈ 1.3) at the first segment, matching those at the HA segment of H3N2 population (Bhatt *et al*. 2011; Kim and Kim 2016).

All three metrics change monotonically (increase in RFD and decrease in λ and TBLD) with increasing reassortment rate, particularly in the range where *r* is between 10^−3^ and 10^−2^ (*N*_e_*r ≅* 0.1 ~ 1) (Figure 2). RFD responds most sensitively to *r*: the distribution of RFD for a given *r* is relatively narrow compared to the change of mean with increasing *r*. However, the absolute values of RFD changes significantly depending on the evolutionary models in the simulation. On the other hand, λ and TBLD exhibit larger variances but are less sensitive to evolutionary models.

**Figure 2.**
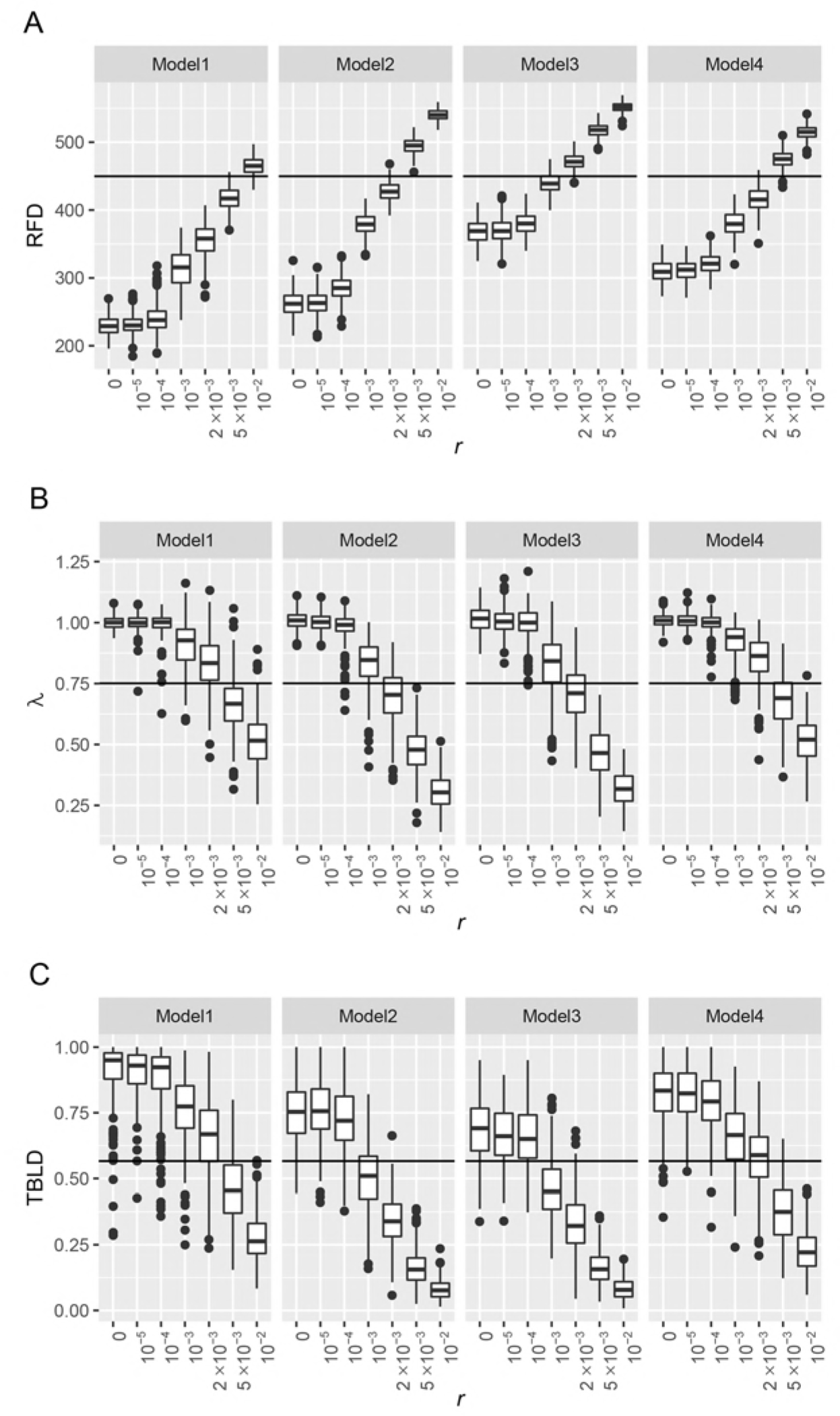
Figure legend: Summary statistics of tree incongruity or linkage disequilibrium with varying reassortment rate (*r*) of H3N2 in simulations. Boxplot of estimates of (A) Robinson-Foulds metric, (B) ratio of between-segment *r*^2^ to within-segment *r*^2^ (λ) and (C) topology-based LD. Each simulation of evolutionary scenario and reassortment rate is run for 300 replicates. A solid line in each plot indicates the average of estimates from HA-PB2, HA-PB1, HA-PA and HA-NP.

We calculated these three metrics from HA-PB2, HA-PB1, HA-PA and HA-NP segment pairs in influenza H3N2 (Table 1). We do not observe clear difference in reassortment rates among these segment pairs. For example, TBLD is smallest between HA and PA but λ is largest for this pair. We therefore take averages over segment pairs and compare them to the simulation results above (see horizontal lines in Figure 2). The agreement between observation and simulation is generally poor for *r* < 10^−3^ or *r* = 10^−2^. Within the range between 10^−3^ and 10^−2^, the most likely value of *r* (judged by difference between the empirical value and the mean of simulated distribution) depends on the combination of metric and simulation model. This may suggest that relationship between each metric and reassortment rate varies according to the pattern of sequence polymorphism in the population or, equivalently, the topology of evolutionary trees shaped by selection and population structure.

**Table 1.** Correlation/incongruence measures for H3N2 viral segment pairs

| segment pair | RFD | $\lambda$ | TBLD | GiRaF<br>(2003-2012) | GiRaF<br>(2007-2016) |
| --- | --- | --- | --- | --- | --- |
| HA-PB2 | 446 | 0.740 | 0.537 | 6 | 3 |
| HA-PB1 | 445 | 0.771 | 0.666 | 8 | 6 |
| HA-PA | 452 | 0.830 | 0.446 | 6 | 5 |
| HA-NP | 456 | 0.665 | 0.617 | 5 | 5 |
| average | 449.75 | 0.751 | 0.566 | 6.25 | 4.75 |

A well-known summary statistic for tree topology is Tajima’s *D* (Tajima 1983). We computed Tajima’s *D*, modified for longitudinally sampled sequences (see Methods), for each segment in actual and simulated data (Table 2, Table S1). Simulation under model 4 yields the values of Tajima’s *D* that closely match the value from the HA segment of H3N2. Therefore, given the hypothesis that tree topology modulates the response of our metrics to reassortment rate, the estimates of *r* under model 4 might be more sensible than under other models. In this case, based on RFD *r* is estimated to be between 0.002 and 0.005. However, it is not clear yet whether Tajima’s *D* captures the aspect of tree topology that modulate the outcome of reassortment or it is tree topology alone that matters. For instance, the relationship between *r* and RFD is quite different in models 2 and 3, which however yield similar values of Tajima’s *D* (Table S1).

**Table 2.**
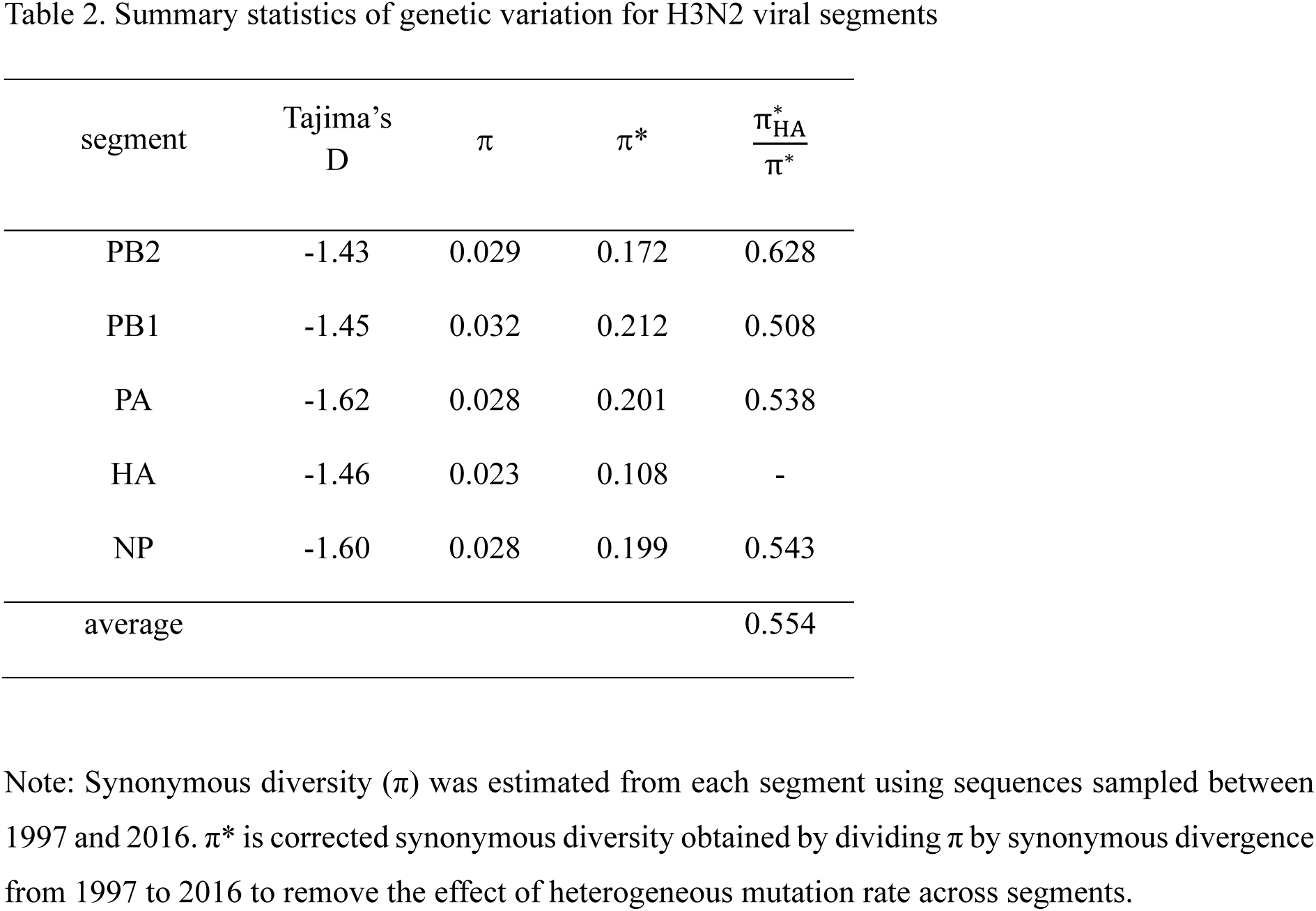
Summary statistics of genetic variation for H3N2 viral segments

| segment | Tajima's<br>D | $\pi$ | $\pi^*$ | $\frac{\pi_{HA}^*}{\pi^*}$ |
| --- | --- | --- | --- | --- |
| PB2 | -1.43 | 0.029 | 0.172 | 0.628 |
| PB1 | -1.45 | 0.032 | 0.212 | 0.508 |
| PA | -1.62 | 0.028 | 0.201 | 0.538 |
| HA | -1.46 | 0.023 | 0.108 | - |
| NP | -1.60 | 0.028 | 0.199 | 0.543 |
| average |  |  |  | 0.554 |

Note: Synonymous diversity ( $\pi$ ) was estimated from each segment using sequences sampled between 1997 and 2016. $\pi^*$ is corrected synonymous diversity obtained by dividing $\pi$ by synonymous divergence from 1997 to 2016 to remove the effect of heterogeneous mutation rate across segments.

Given that *r* is at least 0.002 under the assumption of 80 generations per year, there are approximately 1 − (1 − 0.002)^80^ ≈ 0.15 reassortments per year per viral lineage: namely, one copy of the HA segment and one copy of a non-HA segment found in one virus trace back to two different ancestral viruses of the previous year with more than 15% chance. We confirmed that this per-year estimate does not change when one generation is given 1/160 or 1/40 year (Figure S1).

We next examine how well our inference on reassortment rate matches the result of widely-used method of identifying reassortment events through phylogenetic graph-mining. Using GiRaF (Graph-incompatibility-based Reassortment Finder; Nagarajan and Kingsford 2010), we obtained the candidate sets of reassorted taxa when phylogenies are compared in HA-PB2, HA-PB1, HA-PA, and HA-NP segment pairs (Table 1) and between two segments in the above simulation (Table 3). Simulations show that the numbers of detected reassortments vary greatly according to evolutionary model. Models 2 and 3 lead to a larger number of detection for a given value of *r*. This might be because single-branch reassortments are more detectable with GiRaF (Nagarajan and Kingsford 2010) and genealogies produced under these models have longer outer branches (thus more negative Tajima’s D). With models 1 and 4 (2 and 3), simulation with *r* = 10^−3^ (smaller than 10^−3^) leads to the number of detections similar to that observed in actual viral sequences. Therefore, based on the number of GiRaF-detected reassortment events as a summary statistic, a smaller estimate of *r* is inferred relative to that obtained above using RFD, λ, or TBLD. It however needs to be investigated whether simulated sequences generated under idealized models have allowed better phylogenetic inference and thus more sensitive detection of reassortments.

**Table 3.** The number of candidates sets of reassorted taxa (GiRaF-detected reassortment events) in simulated data.

| $r$ | Model 1 | Model 2 | Model 3 | Model 4 |
| --- | --- | --- | --- | --- |
| $10^{-4}$ | 0.45 | 1.25 | 1.31 | 0.62 |
| $10^{-3}$ | 4.75 | 12.76 | 12.01 | 4.74 |
| $2 \times 10^{-3}$ | 10.1 | 24.71 | 23.62 | 11.03 |
| $5 \times 10^{-3}$ | 22.94 | 49.27 | 26.56 | 16.97 |
| $10^{-2}$ | 38.81 | 69.84 | 71.69 | 40.11 |

Conversely, reassortment rate *r* can be translated into the number of reassortment events on the phylogeny of sampled sequences, which GiRaF targets. Focusing on a specific pair of segments, probability that at least one of two randomly sampled viruses is a reassortant (namely, going backward in time one viral lineage experience reassortment before the coalescence of two lineages occurs) is given approximately by *P*^(2)^ ≡ 2*r*/(1/*N*_e_ + 2*r*) = 2*N*_e_*r*/(2*N*_e_*r* + 1). Then, assuming *r* = 0.001 and *N*_e_ = 140, *P*^(2)^ is about 0.22. With *r* = 0.005 it is about 0.58. Recently, using GiRaF, Berry *et al*. (2016) estimated that about 40% of H3N2 sequences are reassortants, looking at all eight segments. Therefore, the probability of sampling at least one reassortant out of two is 1 − 0.6^2^ ≈ 0.64. Assuming that a random set of segments are exchanged at a reassortment event, this corresponds approximately to *P*^(2)^ = 0.64/2 = 0.32. Considering that GiRaF cannot detect all reassortment events in the sampled genealogy (Nagarajan and Kingsford 2010), we may conclude from this result that the number of reassortment events detected by direct identification of incongruent tree branches is compatible with our estimate of *r* being on the order of 0.001 (~ 0.1 per year).

### Reassortment rate and the inter-segmental pattern of sequence diversity and divergence

Reassortment is critical in shaping the genomic pattern of genetic variation under the effect of selective sweeps and background selection. The more frequent reassortment is, the smaller neutral genetic variation is on a segment under positive selection relative to other segments evolving in more neutral manner. We investigate what range of reassortment rate is compatible with the relative levels of neutral sequence diversity in HA vs. non-HA segments of H3N2. We first calculated synonymous sequence diversities (mean pairwise synonymous differences; π) at the HA, PB2, PB1, PA and NP segments of H3N2 viruses, using sequences sampled from 1997 to 2016 (Table 2). These values however cannot be simply compared with each other because, if mutation rates are not uniform over segments, it can also contribute to differences in neutral genetic diversity. Note that the RNA segments of influenza virus may replicate independently within a host and thus can accumulate mutations at different rates.

Inter-segmental heterogeneity in mutation rate can be detected by differences in synonymous sequence divergence over time. We observe that synonymous substitutions from 1973 to 2016 occur at constant rates at respective segments (Figure 3), in remarkable agreement with molecular clock despite uncertainty regarding whether synonymous substitutions in influenza viruses are strictly neutral or the total number of replication per year is constant over different flu seasons or years. At the same time we note that the rates are variable across segments. For a given pair of segments, the significance of differences in synonymous divergences was evaluated by bootstrapping (Table 4). We find that divergences at HA and PB1 segments, calculated either from four-fold synonymous sites only (Figure 3A) or from all synonymous sites according to Nei-Gojobori method (Figure 3B), are significantly larger than other segments. One may question whether frequent nonsynonymous substitutions in the HA segment have an (unknown) effect of elevating the rate of synonymous substitutions at the same or nearby codons. To test this possibility we measured the synonymous divergences of HA1- and HA2-domain sequences separately. Unlike HA1, on which the epitope sites of hemagglutinin are located, HA2 domain is mainly under stabilizing selection similar to other non-HA genes (Bhatt *et al*. 2011). Synonymous divergence at HA2, obtained from either 4-fold degenerate sites only or using Nei-Gojobori method, is actually larger than that of HA1, although the difference is not significant in the bootstrap test (*p* = 0.16). Therefore, we may rule out the possibility that recurrent nonsynonymous substitutions at HA elevate the rates of synonymous mutations at the corresponding codons.

**Figure 3.**
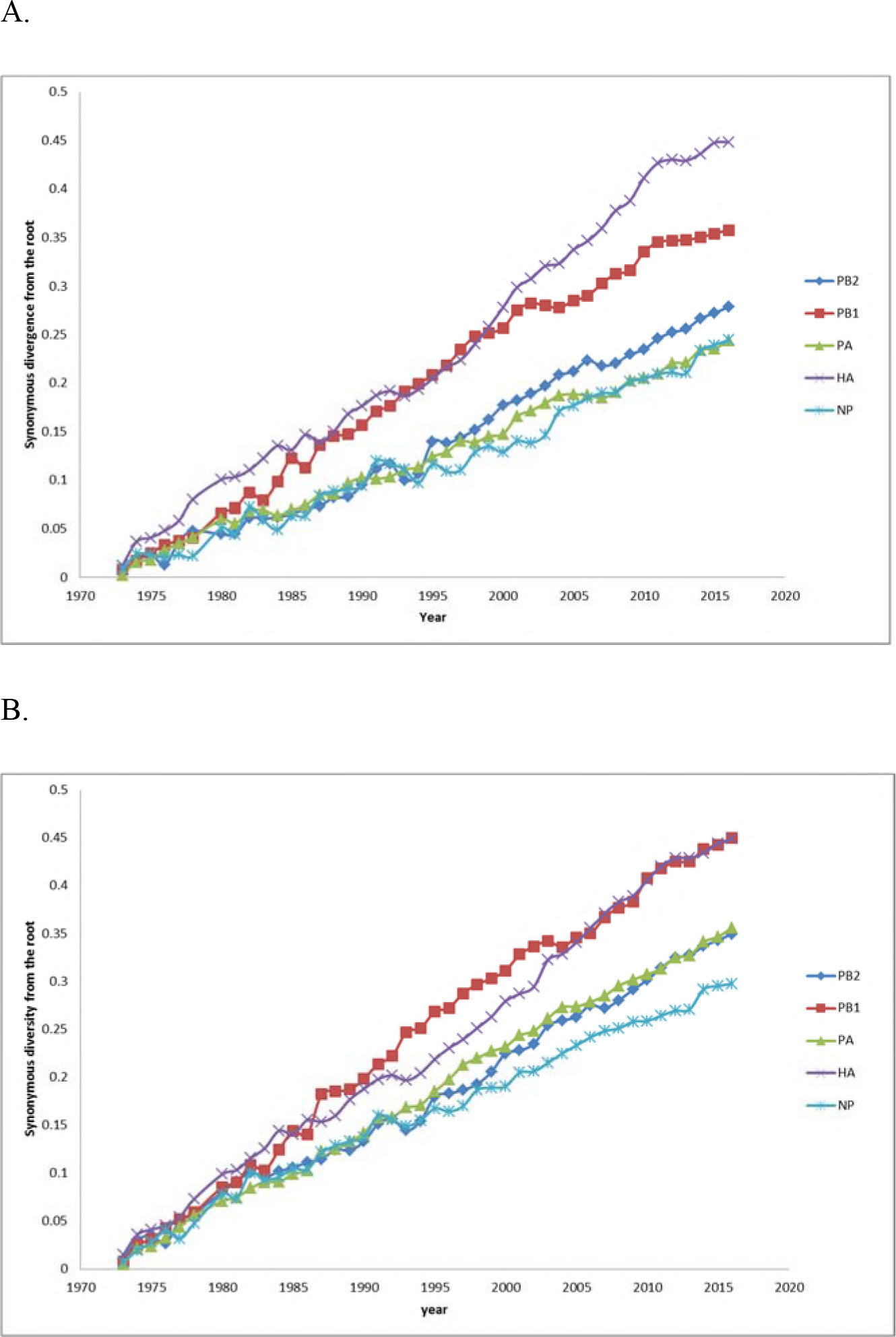
Figure legend: Divergence of H3N2 segments from 1973. Nucleotide divergence from 1973 to each year is calculated from (A) four-fold synonymous sites and (B) synonymous sites according to Neigh-Gojobori method.

**Table 4.** Bootstrap test for heterogenous divergence rate

| Segment | Synonymous sites |  |  |  | Fourfold synonymous sites |  |  |  |
| --- | --- | --- | --- | --- | --- | --- | --- | --- |
|  | PB2 | PB1 | PA | HA | PB2 | PB1 | PA | HA |
| PB1 | 0 |  |  |  | 0 |  |  |  |
| PA | 288 | 20 |  |  | 299 | 8 |  |  |
| HA | 0 | 320 | 0 |  | 0 | 0 | 0 |  |
| NP | 91 | 0 | 42 | 1 | 174 | 5 | 357 | 1 |

Note: The number in each cell indicates the number of bootstrap replicates satisfying $\hat{\theta}^* - \hat{\theta} > \hat{\theta}$ , where $\hat{\theta}$ is the estimated value of difference of divergence rate between two segments and $\hat{\theta}^*$ is the value of $\hat{\theta}$ computed from each bootstrap sample. Bootstrap resampled for 1000 times.

The level of neutral sequence diversity correcting for mutation rate heterogeneity, denoted as π*, is then obtained by dividing π at each segment by its synonymous divergence between 1997 and 2016. π* of HA is about half the level of other segments (Table 2). Synonymous diversity (π) of HA before correction is already lower than those of other segments but the difference becomes larger after the correction. Differences in π* among non-HA segments are small. This result confirms the concentration of positive selection causing selective sweeps on the HA segment.

We next examined which value of *r* best explains the ratio of π* on HA versus non-HA segments. In simulations described above, we calculated neutral diversity on segment 1 and 2, π_1_ and π_2_, and obtained their ratio (π_1_/π_2_) (Figure 4). Since mutation rate is constant in simulation, diversity needs no correction by divergence. We find that reassortment rate close to (in models 2 and 3) or larger than (in model 4) 0.01 best explains the HA vs. non-HA ratio of π* for both *s* = 0.1 (Figure 4) and *s* = 0.05 (Figure S2) This result is not dependent on the frequency of positive selection that we vary to yield different *k*, the number of advantageous substitution per year (Figure S3). Note that the estimate of *r* using correlation statistics (RFD, λ, and TBLD) above is smaller than 0.01. Why a higher rate of reassortment in selective sweep simulations, particularly for model 4, is compatible with π_HA_/π_non-HA_ needs explanation (see Discussion).

**Figure 4.**
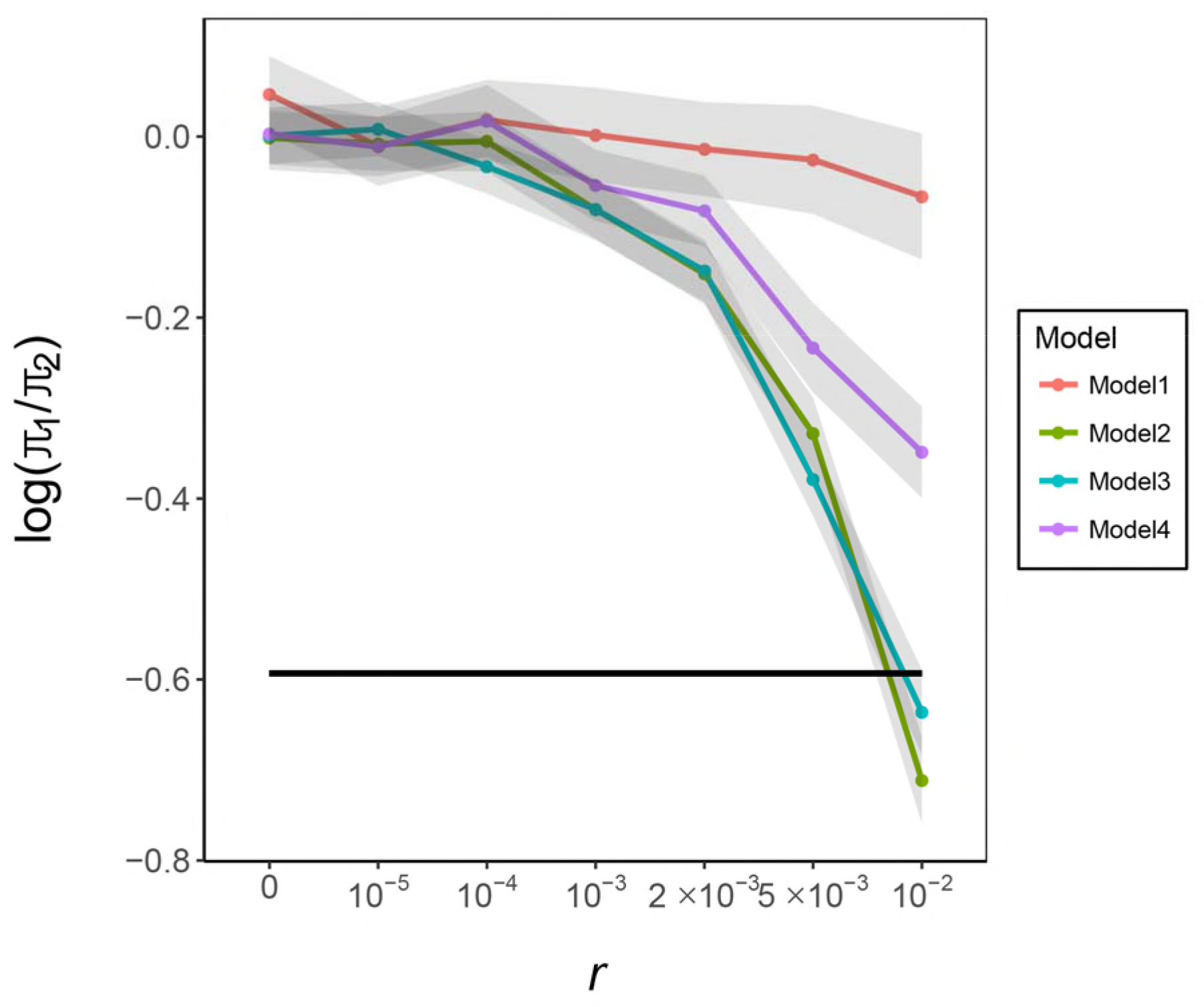
Figure legend: Synonymous diversity of segment 1 relative to segment 2 (π_1_/π_2_) in simulations under different rates of reassortment. The simulated segment 1 mimics HA segment, which is under recurrent positive selection, and the simulated segment 2 mimics a non-antigenic segment (PB2, PB1, PA or NP) of H3N2. Evolutionary models with selection (models 2, 3, and 4) uses *s* = 0.1. A solid horizontal line indicates the observed ratio of π* at HA to mean π* at non-antigenic segments. Gray shades indicate confidential interval given by mean ± 2 standard errors. Note that parameters of each model were adjusted to yield nearly constant π_1_ over values of *r*. Therefore, it is π_2_ that increases with increasing *r*.

To gain further insight on the above result and the dynamics of recurrent selective sweeps, we sought a simple analytic approximation to π_1_/π_2_ using the following heuristic argument. Consider model 2 in which positive selection occurs recurrently in segment 1, generating an equilibrium flux of beneficial alleles reaching fixation in a single constant-sized population. Discrete events of sweeps can be arranged in order, backward in time: let allele *B*_1_ be a beneficial allele that was fixed in the last sweep. (There can be multiple beneficial alleles at different sites being fixed together at a single episode of sweep due to temporal clustering of substitutions (Kim and Kim 2015). In that case, *B*_1_ represents the one that originated by most recent mutation.) The beneficial alleles fixed in the preceding rounds of sweeps are defined as *B*_2_, *B*_3_, and so on. The allele frequency of *B_i_* is given by *x_i_*. Two randomly chosen copies of segment 1 have their most recent common ancestor at *t*_1_ generations back in time. We may assume that *t*_1_ is distributed with mean τ_1_ that is determined only by the rate of selective sweeps. Namely, coalescence due to genetic drift during time interval between successive sweeps is ignored. Then, tracing events backward in time from present, coalescence occurs as *x*_1_ approaches close to zero. Therefore *t*_1_ should be slightly smaller than waiting time until the time of *B*_1_’s entrance into the population. It is also possible that, at the time of sampling lineages, there is a currently sweeping beneficial allele that has not reached fixation. If both sampled lineages carry this sweeping allele, their coalescence should occur close to the time of this beneficial allele’s entrance. Here we simply define τ_1_ as the mean of *t*_1_ when all of such possibilities under the equilibrium flux of beneficial alleles are taken into account.

With complete linkage (*r* = 0), identical backward-in-time process governs the coalescence of randomly sampled lineages in segment 2 and their mean coalescent time is τ_1_. However, with *r* > 0 two lineages may avoid coalescence by reassortment: at a given generation, each lineage can recombine away from B1 allele with probability *r*(1 - *x*_1_). Given that *r*/*s* ≪ 1, where *s* is the strength of selection, the probability of such a lineage recombining back to *B*_1_ is very small and thus can be ignored. The opportunity for a lineage to recombine away increases as *x*_1_ remains longer at low values (but not too low forcing coalescence). Therefore, the length of trajectory *x*_1_ determines the probability of escaping coalescence. While *x*_1_ should increase from 1/*N* to 1, forward-in-time, stochastic effect makes the trajectory much shorter than the length of deterministic trajectory: the change of *x*_1_ is approximated by instantaneous increase from 1/*N* to 1/(*Nf*), where *f* is the fixation probability of a copy of beneficial allele (Maynard Smith 1971), followed by deterministic increase expected for selective advantage *s*. Then, using the approximation obtained in (Barton 2000) and other studies, the probability of escaping coalescence in one round of sweep is given by

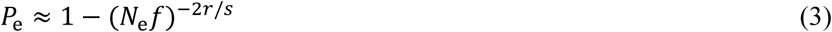

where *N*_e_ is the effective population size under which sweeps occur (i.e, *N*_e1_ of (Kim and Kim 2016)). Now, two lineages that have just escaped coalescence are subject to coalescence in the next (earlier) round of sweep by *B*_2_. Assuming that successive sweeps occur as a random Poisson process, the waiting time until *x*_2_ becomes small enough to force coalescence is again τ_1_. Then, if the lineages coalesce in the *n*^th^ round of sweep, it takes on average *nτ*_1_ generations. Therefore, the mean coalescent time for segment 2 is

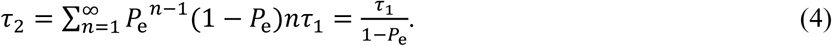

The level of sequence diversity on segment 1 relative to that on segment 2 is therefore

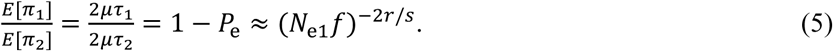

This approximation shows that π_1_/π_2_ does not depend on the rate of recurrent positive selection (*k*) but on the strength of selection, in agreement with our simulation (Figure S3).

We compared the simulation results of model 2 with Eq. (5) in which *f* is replaced by either 2*s*, a usual approximation under infrequent selective sweeps, or mean fixation probability observed in simulation. The latter is 0.0269 for *s* = 0.1 and 0.0253 for *s* = 0.05. Therefore, actual fixation probabilities are much smaller than 2*s*, indicating that strong clonal interference occurs in our simulated populations (i.e. under parameters constrained to yield both π_1_ ≈ 0.027 per site and *k* ≈ 1.3 per year). Figure 5 shows that π_1_/π_2_ predicted by eq. (5), using either choices of *f*, is much smaller than that observed in simulation. Namely, lineages on segment 2 in simulation do not escape coalescence as frequently as predicted under the assumption of eq. (5), producing π_2_ not so larger than π_1_. It might be suggested that, in addition to the initial acceleration of *x_i_* by a factor of 1/*f*, *x_i_* would increase much faster than expected with selection coefficient *s* under clonal interference, because successful beneficial mutations reaching fixation tend to form temporal clusters, i.e. in positive linkage disequilibrium with each other (Strelkowa and Lassig 2012; Kim and Kim 2015). However, when we estimated the “effective” selection coefficients of beneficial alleles by counting generations that the sample frequency of *x_i_* takes to increase from ~0.2 to ~0.8 for all trajectories in simulation with *s* = 0.1, the mean was 0.069. We therefore did not obtain an evidence of faster increase in *x_i_* by clonal interference. It remains to be investigated what causes coalescence to occur faster, relative to recombination, than expected by eq. (5).

**Figure 5.**
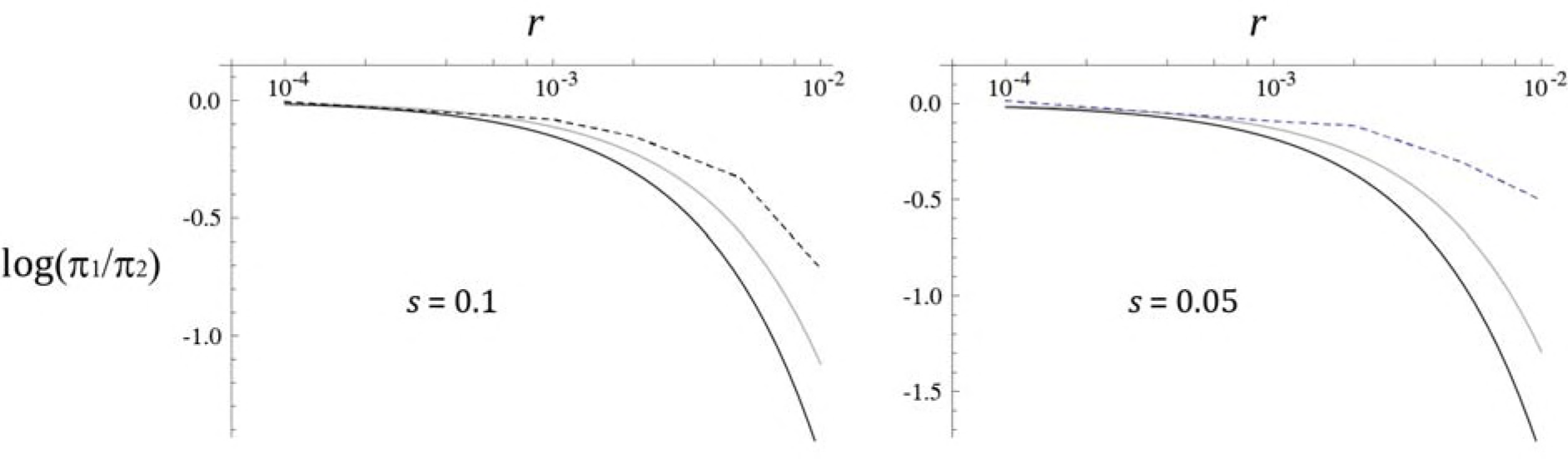
Figure legend: Synonymous diversity of segment 1 relative to segment 2 (π_1_/ π_2_) in simulations under different rates of reassortment (*r*) predicted by eq. (5) using *f* = 2*s* (dark curve) or using the observed fixation probability (0.0269 for *s* = 0.1 and 0.0253 for *s* = 0.05) for *f*. Simulation results for model 2 are shown by points connected by dashed lines.

## DISCUSSION

The population genetics of sexually reproducing organisms demonstrated that recombination rate is as important as other fundamental evolutionary parameters, such as mutation rate, effective population size, and selection coefficient, for understanding their evolution and genetic diversity. Therefore, in order to build a correct model that predicts the direction of influenza viral evolution, the rate of reassortment relative to other parameters needs to be estimated. While reassortment occurs in all 28 pairs of influenza viral segments, this study focused on reassortment between the HA segment, the major target of strong positive selection, and one of four non-HA segments (PB2, PB1, PA, NP) that are generally considered to evolve neutrally (Bhatt *et al*. 2011), because such reassortment is expected to cause difference in inter-segmental difference in genetic diversity and is therefore key to inferring positive selection in influenza virus. The NA (neuraminidase) segment was not included among non-HA segments because, with epistatic interactions detected between HA and NA genes (Neverov *et al*. 2014), particular reassortants may have higher or lower fitness relative to non-reassortants and can thus bias the estimate of reassortment rate. MP and NS segments, also known to undergo little positive selection, were not included because their synonymous sites are not likely to evolve neutrally due to overlapping protein-coding regions. We used multiple summary statistics of correlation or congruence between segmental sequence diversity to infer the range of reassortment rate in the H3N2 viral population. In general, it is suggested that the probability of reassortment per virus per viral infection cycle is between 0.001 and 0.01. Ideally, information from multiple summary statistics might be combined to yield a narrower range of estimate for example using approximate Bayesian computation (ABC) (Beaumont *et al*. 2002). However, our individual-based simulation was too slow for such implementation. It might be possible in the future to develop a coalescent-based simulation that is fast enough for ABC. Since relationships between the summary statistics and reassortment rate depends on the evolutionary model of virus, such approach will have to estimate reassortment rate jointly with other parameters of selection and demography.

Reassortment rate determines the hitchhiking effect of recurrent positive selection at the HA segment on neutral genetic variation at other segments (Figure 4). Our simulation results suggest that more frequent reassortment (*r* ≥ 0.01) than inferred above using correlation statistics (i.e. 0.001 ≤ *r* < 0.01) is needed to explain ~40% lower synonymous diversity on HA relative to those on non-HA segments, particularly under complex demography (model 4). Given that our earlier inference of *r* < 0.01 is correct, this discrepancy would indicate that, for a given *r*, neutral lineages on non-HA segments escape hitchhiking (i.e. avoid coalescence forced by a sweep) more frequently than expected under the simulation models. It might be because our simulation models use reassortment rates that are constant in the course of selective and demographic dynamics (even in model 4). Namely, it assumes that hosts experience a constant rate of coinfection through time. This is not likely true in the H3N2 population, in which coinfection must be more frequent during the seasonal peaks of population size (the number of hosts infected). Then, if a new immunity-evading adaptive allele is more likely to arise during peaks of influenza epidemics, as expected from the principle that mutational input is proportional to population size and the fixation probability of adaptive allele is larger during the period of population size expansion (Otto and Whitlock 1997), this adaptive allele may be transmitted through coinfected hosts more frequently, thus participating in more reassortment, than non-adaptive alleles. Therefore, because hitchhiking effect is mostly determined during the early phase when the adaptive variant is still in low frequency (Maynard Smith and Haigh 1974), the observed ratio of HA to non-HA synonymous diversity can be explained by an effectively higher reassortment rate experienced by antigenic variants on the HA segment. Further investigation on adaptive evolution and inter-segmental diversity in influenza virus will require a theoretical/simulation model that allows realistic seasonal influenza dynamics and associated change in coinfection/reassortment rate.

Wider discrepancy between data and the model of positive selection under metapopulation structure (model 4 in Figure 4) demands further theoretical explanations. At first, it is known that the spatial structure of a population slows down the spread of a beneficial allele across demes, thus weakening the hitchhiking effect as there are more opportunities for neutral lineages to recombine away from the beneficial allele (Kim and Maruki 2011; Barton *et al*. 2013). This contradicts with our result: if hitchhiking effect is weaker, π_1_/π_2_ should become smaller for a given *r*. However, demographic model assumed in those studies are quite different from the one used here. In our model 4, seven out of eight demes undergo extinction-recolonization cycles. While a beneficial allele is increasing in frequency in the total population, “empty” demes are more likely to be colonized by viruses carrying this than the ancestral allele. (Note that our model assigns the absolute fitness to haploid individuals so that a local population can be established even from a single immigrant (Kim and Kim 2016).) Because no or small number of individuals carrying the non-beneficial allele exist where those carrying beneficial allele increase exponentially, neutral lineages on segment 2 can hardly escape coalescence, thus resulting in stronger reduction in polymorphism. This stronger hitchhiking effect during the establishment of a new local population was demonstrated in the model of “Genotype-Dependent Colonization and Introgression (GDCI)” in Kim and Gulisija (2010). Unless *r* is very larger than 0.01 (or the joint effect of selection and co-infection dynamics increasing the effective recombination rate considered above is very dramatic), the overestimation of π_1_/π_2_ by model 4 may suggest that selective sweeps in the actual population of H3N2 do not occur predominantly through GDCI process. While the transmission of influenza virus in most regions of northern and southern hemispheres is seasonal, continuous year-round transmission occurs in certain tropical or subtropical regions (Viboud *et al*. 2006). Selective sweeps in such continuous viral populations would not involve the GDCI process. Therefore, if global influenza genetic diversity is mainly shaped by variants arising from the permanent tropical populations (Rambaut *et al*. 2008; Chan *et al*. 2010), the overall effects of selective sweeps might be closer to those in our models 2 and 3. In our simulation of model 4, one out of eight demes are maintained at a constant size. Its small size however might have limited its contribution to the diversity of the total population.

While not initially a major focus of this study, significant inter-segmental heterogeneity in the rate of synonymous substitutions, indicating that new mutations occur at different rates in different segments, is an unexpected discovery. Negative-sense viral RNA strands replicate via positive-sense mRNA strands. Then, if segments are transcribed at different rates for example due to different demands for or turn-over rates of viral proteins, some segments may experience more negative-positive-negative replication cycles than others before being assembled into viral particles. Such difference would translate into different mutation rates given the fixed rate of RNA replication errors per cycle. We may also speculate that replication error is influenced by the secondary structures of RNA strands that are probably different among segments.

## ACKNOWLEDGEMENT

This research was supported by the National Research Foundation of Korea grants 2012R1A1A2004932 to YK.

## SUPPORTING INFORMATION

**Table S1.**
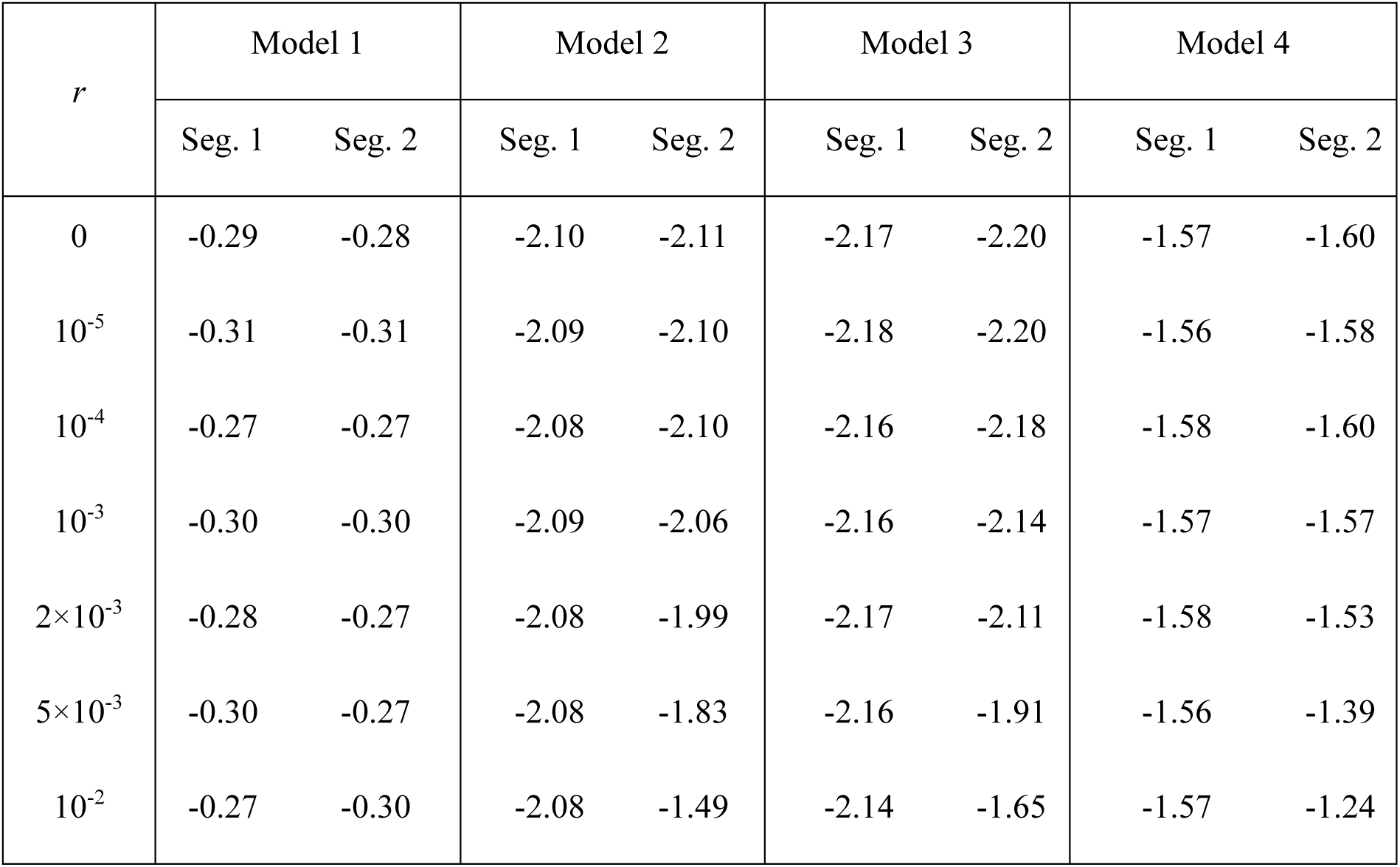
Average Tajima’s D in simulations

| $r$ | Model 1 | | Model 2 | | Model 3 | | Model 4 | |
| --- | --- | --- | --- | --- | --- | --- | --- | --- |
|  | Seg. 1 | Seg. 2 | Seg. 1 | Seg. 2 | Seg. 1 | Seg. 2 | Seg. 1 | Seg. 2 |
| 0 | -0.29 | -0.28 | -2.10 | -2.11 | -2.17 | -2.20 | -1.57 | -1.60 |
| $10^{-5}$ | -0.31 | -0.31 | -2.09 | -2.10 | -2.18 | -2.20 | -1.56 | -1.58 |
| $10^{-4}$ | -0.27 | -0.27 | -2.08 | -2.10 | -2.16 | -2.18 | -1.58 | -1.60 |
| $10^{-3}$ | -0.30 | -0.30 | -2.09 | -2.06 | -2.16 | -2.14 | -1.57 | -1.57 |
| $2 \times 10^{-3}$ | -0.28 | -0.27 | -2.08 | -1.99 | -2.17 | -2.11 | -1.58 | -1.53 |
| $5 \times 10^{-3}$ | -0.30 | -0.27 | -2.08 | -1.83 | -2.16 | -1.91 | -1.56 | -1.39 |
| $10^{-2}$ | -0.27 | -0.30 | -2.08 | -1.49 | -2.14 | -1.65 | -1.57 | -1.24 |

**Figure S1.**
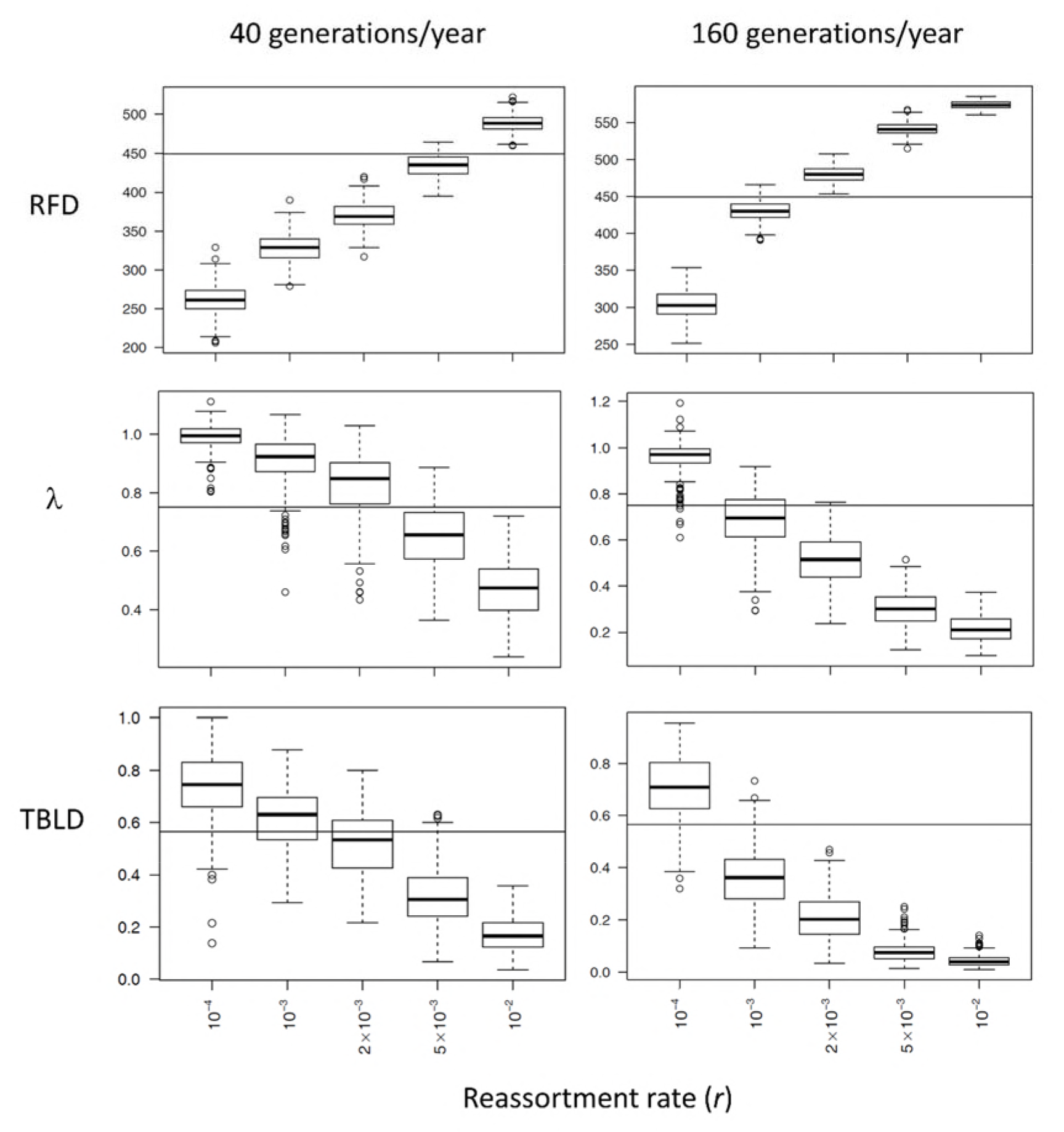
Figure S1 legend: Correlation/incongruence statistics in simulations of model 2 with varying reassortment rate when the number of generations is 40 or 160 per year. Four-fold increase in generations/year led to reduction in the estimates of *r* (per generation) by approximately the same factor, yielding approximately constant reassortment rate per year. Note that, relative to simulation with 80 generations per year, population size decreases (increases) and mutation rate/generation increases (decreases) by a factor of ~2 in the simulation with 40 (160) generations/year to produce the equivalent level of synonymous polymorphism.

**Figure S2.**
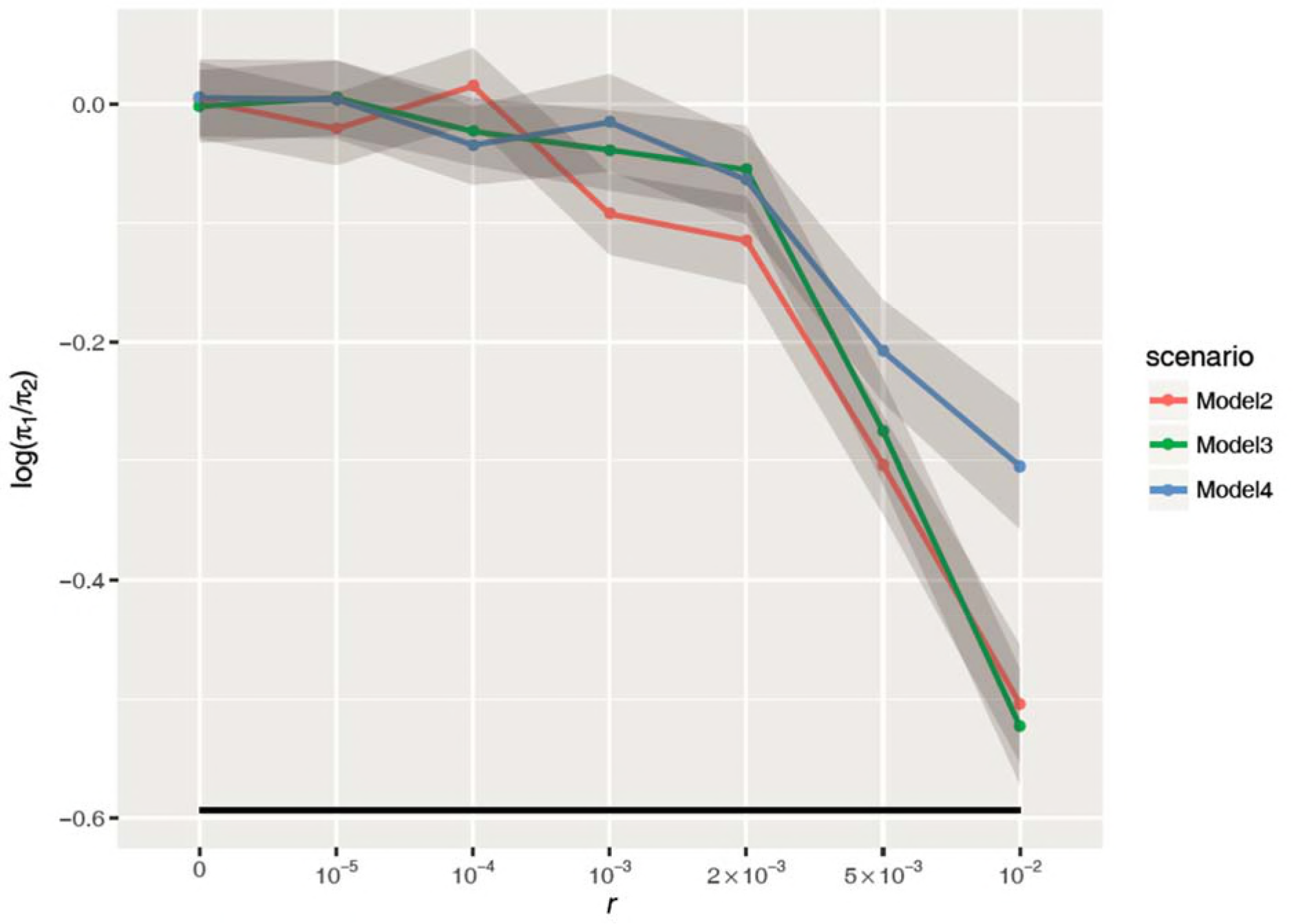
Figure S2 legend: Synonymous diversity of segment 1 relative to segment 2 (π_1_/ π_2_) in simulations under different rates of reassortment with *s* = 0.05. A solid horizontal line indicates the observed ratio of π* at HA to mean π* at non-antigenic segments. Gray shades indicate confidential interval given by mean ± 2 standard errors.

**Figure S3.**
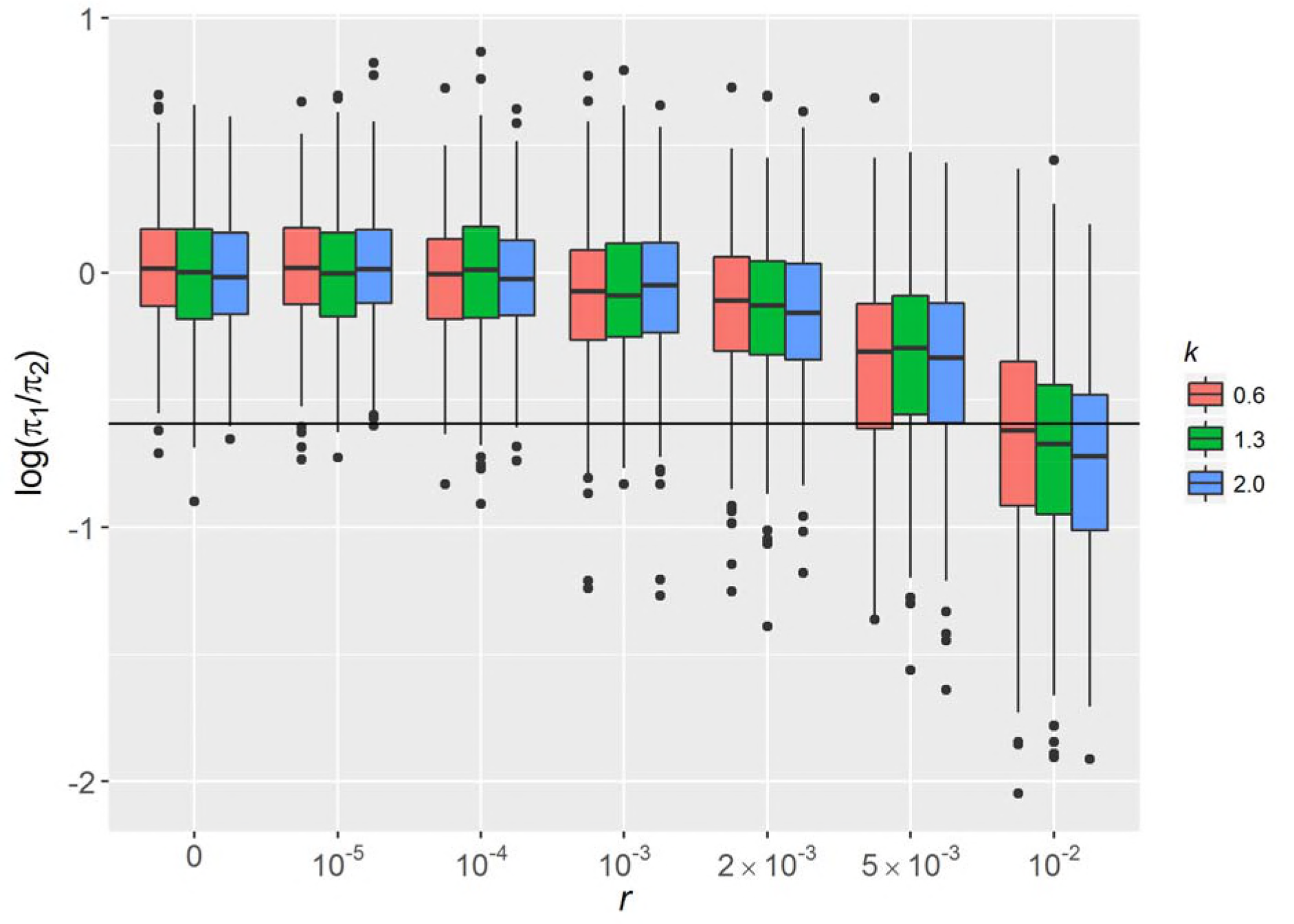
Figure S3 legend: Synonymous diversity of segment 1 relative to segment 2 (π_1_/ π_2_) in simulations with different reassortment rates (*r*) and adaptive substitution rates (*k*). Results of model 2 only are shown.

## LITERATURE CITED

Barton, N. H., 2000 Genetic hitchhiking. Philosophical Transactions of the Royal Society B: Biological Sciences 355: 1553–1562.

Barton, N. H., A. M. Etheridge, J. Kelleher and A. Véber, 2013 Genetic hitchhiking in spatially extended populations. Theoretical population biology 87: 75–89.

Beaumont, M. A., W. Zhang and D. J. Balding, 2002 Approximate Bayesian Computation in Population Genetics. Genetics 162: 2025–2035.

Bedford, T., S. Cobey, P. Beerli and M. Pascual, 2010 Global migration dynamics underlie evolution and persistence of human influenza A (H3N2). PLoS pathogens 6: e1000918.

Bedford, T., S. Cobey and M. Pascual, 2011 Strength and tempo of selection revealed in viral gene genealogies. BMC Evolutionary Biology 11: 220.

Berry, I. M., M. C. Melendrez, T. Li, A. W. Hawksworth, G. T. Brice et al., 2016 Frequency of influenza H3N2 intra-subtype reassortment: attributes and implications of reassortant spread. BMC biology 14: 117.

Bhatt, S., E. C. Holmes and O. G. Pybus, 2011 The genomic rate of molecular adaptation of the human influenza A virus. Mol Biol Evol 28: 2443–2451.

Buonagurio, D. A., S. Nakada, J. D. Parvin, M. Krystal, P. Palese et al., 1986 Evolution of human influenza A viruses over 50 years: rapid, uniform rate of change in NS gene. Science 232: 980–982.

Chan, J., A. Holmes and R. Rabadan, 2010 Network analysis of global influenza spread. PLoS Comput Biol 6: e1001005.

Charlesworth, B., M. T. Morgan and D. Charlesworth, 1993 The effect of deleterious mutations on neutral molecular variation. Genetics 134: 1289–1303.

Dudas, G., T. Bedford, S. Lycett and A. Rambaut, 2014 Reassortment between influenza B lineages and the emergence of a coadapted PB1–PB2–HA gene complex. Molecular biology and evolution 32: 162–172.

Ferguson, N. M., A. P. Galvani and R. M. Bush, 2003 Ecological and immunological determinants of influenza evolution. Nature 422: 428–433.

Fitch, W. M., R. M. Bush, C. A. Bender and N. J. Cox, 1997 Long term trends in the evolution of H(3) HA1 human influenza type A. Proc Natl Acad Sci U S A 94: 7712–7718.

Fitch, W. M., J. M. Leiter, X. Q. Li and P. Palese, 1991 Positive Darwinian evolution in human influenza A viruses. Proc Natl Acad Sci U S A 88: 4270–4274.

Hall, P., and S. R. Wilson, 1991 Two guidelines for bootstrap hypothesis testing. Biometrics: 757–762.

Hill, W., and A. Robertson, 1968 Linkage disequilibrium in finite populations. Theoretical and Applied Genetics 38: 226–231.

Holmes, E. C., E. Ghedin, N. Miller, J. Taylor, Y. Bao et al., 2005 Whole-genome analysis of human influenza A virus reveals multiple persistent lineages and reassortment among recent H3N2 viruses. PLoS biology 3: e300.

Illingworth, C. J., and V. Mustonen, 2012 Components of selection in the evolution of the influenza virus: linkage effects beat inherent selection. PLoS pathogens 8: e1003091.

Ina, Y., and T. Gojobori, 1994 Statistical analysis of nucleotide sequences of the hemagglutinin gene of human influenza A viruses. Proc Natl Acad Sci U S A 91: 8388–8392.

Kim, K., and Y. Kim, 2015 Episodic Nucleotide Substitutions in Seasonal Influenza Virus H3N2 Can Be Explained by Stochastic Genealogical Process without Positive Selection. Molecular Biology and Evolution 32: 704–710.

Kim, K., and Y. Kim, 2016 Population genetic processes affecting the mode of selective sweeps and effective population size in influenza virus H3N2. BMC evolutionary biology 16: 156.

Kim, Y., and D. Gulisija, 2010 Signatures of recent directional selection under different models of population expansion during colonization of new selective environments. Genetics 184: 571–585.

Kim, Y., and T. Maruki, 2011 Hitchhiking effect of a beneficial mutation spreading in a subdivided population. Genetics 189: 213–226.

Koel, B. F., D. F. Burke, T. M. Bestebroer, S. van der Vliet, G. C. Zondag et al., 2013 Substitutions near the receptor binding site determine major antigenic change during influenza virus evolution. Science 342: 976–979.

Lu, L., S. J. Lycett and A. J. L. Brown, 2014 Reassortment patterns of avian influenza virus internal segments among different subtypes. BMC evolutionary biology 14: 16.

Lycett, S., G. Baillie, E. Coulter, S. Bhatt, P. Kellam et al., 2012 Estimating reassortment rates in co-circulating Eurasian swine influenza viruses. Journal of General Virology 93: 2326–2336.

Maynard Smith, J., 1971 What use is sex? Journal of theoretical biology 30: 319–335.

Maynard Smith, J., and J. Haigh, 1974 The hitch-hiking effect of a favorable gene. Genet. Res. 23: 23–35.

McDonald, S. M., M. I. Nelson, P. E. Turner and J. T. Patton, 2016 Reassortment in segmented RNA viruses: mechanisms and outcomes. Nature Reviews Microbiology 14: 448.

Nagarajan, N., and C. Kingsford, 2010 GiRaF: robust, computational identification of influenza reassortments via graph mining. Nucleic acids research 39: e34-e34.

Nei, M., and T. Gojobori, 1986 Simple methods for estimating the numbers of synonymous and nonsynonymous nucleotide substitutions. Molecular biology and evolution 3: 418–426.

Nelson, M. I., and E. C. Holmes, 2007 The evolution of epidemic influenza. Nat Rev Genet 8: 196–205.

Neverov, A. D., K. V. Lezhnina, A. S. Kondrashov and G. A. Bazykin, 2014 Intrasubtype reassortments cause adaptive amino acid replacements in H3N2 influenza genes. PLoS genetics 10: e1004037.

Otto, S. P., and M. C. Whitlock, 1997 The probability of fixation in populations of changing size. Genetics 146: 723–733.

Pinsent, A., C. Fraser, N. M. Ferguson and S. Riley, 2015 A systematic review of reported reassortant viral lineages of influenza A. BMC infectious diseases 16: 3.

Pybus, O. G., A. Rambaut, R. Belshaw, R. P. Freckleton, A. J. Drummond et al., 2007 Phylogenetic evidence for deleterious mutation load in RNA viruses and its contribution to viral evolution. Mol Biol Evol 24: 845–852.

Rabadan, R., A. J. Levine and M. Krasnitz, 2008 Non-random reassortment in human influenza A viruses. Influenza and Other Respiratory Viruses 2: 9–22.

Rambaut, A., O. G. Pybus, M. I. Nelson, C. Viboud, J. K. Taubenberger et al., 2008 The genomic and epidemiological dynamics of human influenza A virus. Nature 453: 615–619.

Robinson, D. F., and L. R. Foulds, 1981 Comparison of phylogenetic trees. Mathematical biosciences 53: 131–147.

Schweiger, B., L. Bruns and K. Meixenberger, 2006 Reassortment between human A (H3N2) viruses is an important evolutionary mechanism. Vaccine 24: 6683–6690.

Simonsen, L., C. Viboud, B. T. Grenfell, J. Dushoff, L. Jennings et al., 2007 The genesis and spread of reassortment human influenza A/H3N2 viruses conferring adamantane resistance. Molecular biology and evolution 24: 1811–1820.

Strelkowa, N., and M. Lassig, 2012 Clonal interference in the evolution of influenza. Genetics 192: 671–682.

Swofford, D. L., 2003 PAUP*: phylogenetic analysis using parsimony, version 4.0 b10.

Tajima, F., 1989 Statistical method for testing the neutral mutation hypothesis by DNA polymorphism. Genetics 123: 585–595.

Viboud, C., W. J. Alonso and L. Simonsen, 2006 Influenza in tropical regions. PLoS medicine 3: e89.

Villa, M., and M. Lässig, 2017 Fitness cost of reassortment in human influenza. PLoS pathogens 13: e1006685.

Westgeest, K. B., C. A. Russell, X. Lin, M. I. Spronken, T. M. Bestebroer et al., 2014 Genomewide analysis of reassortment and evolution of human influenza A (H3N2) viruses circulating between 1968 and 2011. Journal of virology 88: 2844–2857.

Wirtz, J., M. Rauscher and T. Wiehe, 2018 Topological linkage disequilibrium calculated from coalescent genealogies. bioRxiv.

Yang, Z., 2000 Maximum likelihood estimation on large phylogenies and analysis of adaptive evolution in human influenza virus A. J Mol Evol 51: 423–432.

